# 3D cell culture models demonstrate a role for FGF and WNT signaling in regulation of lung epithelial cell fate and morphogenesis

**DOI:** 10.1101/2020.04.04.024943

**Authors:** Anas Rabata, Radek Fedr, Karel Soucek, Ales Hampl, Zuzana Koledova

**Affiliations:** Department of Histology and Embryology, Faculty of Medicine, Masaryk University, Kamenice 3, Brno, 625 00, Czech Republic; Department of Cytokinetics, Institute of Biophysics of the Czech Academy of Sciences, Brno, Czech Republic; International Clinical Research Center, St. Anne’s University Hospital Brno, Brno, Czech Republic

**Author notes:** To whom correspondence should be addressed: Zuzana Koledova, Ales Hampl.

**Keywords:** 3D cell culture, epithelial cell, FGF signaling, lung, morphogenesis, organoid, WNT signaling

## Abstract

FGF signaling plays an essential role in lung development, homeostasis, and regeneration. Several FGF ligands were detected in the developing lungs, however, their roles have not been fully elucidated. We employed mouse 3D cell culture models and imaging to *ex vivo* study of *a)* the role of FGF ligands in lung epithelial morphogenesis and *b)* the interplay of FGF signaling with epithelial growth factor (EGF) and WNT signaling pathways. In non-adherent conditions, FGF signaling promoted formation of lungospheres from lung epithelial stem/progenitor cells (LSPCs). Based on their architecture, we defined three distinct phenotypes of lungospheres. Ultrastructural and immunohistochemical analyses showed that LSPCs produced more differentiated lung cell progeny. In 3D extracellular matrix, FGF2, FGF7, FGF9, and FGF10 promoted lung organoid formation with similar efficiency. However, FGF9 showed reduced capacity to promote lung organoid formation, suggesting that FGF9 has a reduced ability to sustain LSPCs survival and/or initial divisions. Analysis of lung organoid phenotypes revealed that FGF7 and FGF10 produce bigger organoids and induce organoid branching with higher frequency than FGF2 and FGF9. Higher FGF concentration and/or the use of FGF2 with increased stability and affinity to FGF receptors both increased lung organoid and lungosphere formation efficiency, respectively, suggesting that the level of FGF signaling is a crucial driver of LSPC survival and differentiation, and also lung epithelial morphogenesis. EGF signaling played a supportive but nonessential role in FGF-induced lung organoid formation. Moreover, analysis of tissue architecture and cell type composition confirmed that the lung organoids contained alveolar-like regions with cells expressing alveolar type I and type II cell markers, as well as airway-like structures with club cells and ciliated cells. WNT signaling enhanced the efficiency of lung organoid formation, but in the absence of FGF10 signaling, the organoids displayed limited branching and less differentiated phenotype. In summary, we present lung 3D cell culture models as useful tools to study the role and interplay of signaling pathways in lung development and we reveal roles for FGF ligands in regulation of mouse lung morphogenesis *ex vivo*.

## Introduction

Mammalian lung is a complex, stereotypically branched organ, whose development is strictly regulated by multiple signaling pathways. They include fibroblast growth factor (FGF) (Lebeche et al., 1999; Ohuchi et al., 2000; Tichelaar et al., 2000; del Moral et al., 2006; Cardoso and Lü, 2006), WNT (Shu et al., 2005a; Yin et al., 2008), bone morphogenetic protein (Weaver et al., 1999; Lu et al., 2001; Shu et al., 2005b), Sonic Hedgehog (Miller et al., 2004; White et al., 2006), epidermal growth factor (EGF) (Kheradmand et al., 2002), retinoic acid (Malpel et al., 2000), and HIPPO (Volckaert et al., 2019) pathways. Tight interplay of these pathways is essential also for lung epithelial homeostasis, regeneration, and repair (Volckaert and De Langhe, 2015; Volckaert et al., 2017).

FGF signaling plays an essential role in lung development from the very earliest stages. FGF4 is involved in organ-specific domain formation along the anteroposterior axis of the early endoderm (Wells and Melton, 2000; Dessimoz et al., 2006). Later, FGF10-FGFR2b signaling is essential for lung bud formation and to control branching morphogenesis of the lung. Deletion of either *Fgf10* or *Fgfr2b* results in complete distal lung agenesis (Min et al., 1998; Sekine et al., 1999; De Moerlooze et al., 2000), while *Fgf10* hypomorphic lungs display decreased ramifications (Ramasamy et al., 2007). *Fgf10* gain-of-function prevents differentiation of epithelial tip cells toward the bronchial progenitor lineage and disrupts lung morphogenesis (Nyeng et al., 2008; Volckaert et al., 2013). Furthermore, FGF1, FGF2, FGF7, and FGF9, were found in fetal rodent lung, too (Han et al., 1992; Cardoso et al., 1997; Powell et al., 1998; Colvin et al., 2001; Jones et al., 2019). FGF7 acts as a proliferative factor for lung epithelium during lung development (Lebeche et al., 1999) and together with FGF2 it induces expression of surfactant proteins (Matsui et al., 1999). FGF9 is responsible for mesenchymal cell proliferation, and it is also involved in lung epithelium regulation (del Moral et al., 2006).

The role of FGF signaling in lung development is interwoven with WNT signaling. FGF and WNT signaling regulate proximal/distal patterning and fate of lung progenitor cells (Volckaert and De Langhe, 2015). Canonical WNT signaling is required for mesenchymal expression of FGF10 and primary lung bud formation (Goss et al., 2009). Furthermore, mesenchymal WNT signaling regulates amplification of *Fgf10* expressing airway smooth muscle cell progenitors in the distal mesenchyme (Volckaert and De Langhe, 2015). In adult lung, FGF10 and WNT signaling regulate the activity of basal cells, the lung epithelial stem/progenitor cells (LSPCs) that ensure lung epithelial homeostasis and repair after injury (Volckaert et al., 2013). However, the exact functions of FGF and WNT signaling in LSPCs have not been fully elucidated.

In this study, we investigated the role of FGF and WNT signaling in regulation of epithelial morphogenesis from LSPCs. To this end, we developed and used several 3D cell culture techniques, including lungosphere and lung organoid assays, and we investigated the ability of various FGF ligands and WNT signaling to support LSPC survival and differentiation to epithelial structures.

## Results

### Lungosphere assay demonstrates the existence of cells with capacity for anchorage-independent growth and self-renewal

Stem and progenitor cells are defined by their capacities to self-renew (i.e. to replicate and form more of the same cells), as well as to produce more differentiated progeny (Fuchs and Chen, 2013). On top of that, one of the distinctive characteristics of stem and progenitor cells is their ability to resist anoikis and to survive in non-adherent conditions (Pastrana et al. 2011). These characteristics have been applied in sphere formation assays, such as neurosphere (Reynolds and Weiss, 1992) or mammosphere (Shaw et al., 2012) formation assays and to some extent also in lung cancer sphere formation assays (Zhao et al. 2015). We applied this approach to isolate LSPCs. Single epithelial cells from mouse lung were seeded in non-adherent plates in defined serum-free medium with epidermal growth factor (EGF) and FGF2 and cultured for 10 to 14 days, with addition of fresh medium every 3 days (Shaw et al., 2012; Rabata et al., 2017). Because FGF2 rapidly loses its biological activity at 37°C, we tested the use of FGF2-wt, as well as FGF2 with increased thermal stability (FGF2-STAB) (Dvorak et al., 2018) and sustained FGFR specificity (Koledova et al., 2019) for their capacity to support lungosphere formation. With FGF2-wt, the lungosphere formation efficiency (LFE) was 0.088 ± 0.006% (Figure S1A, B) and the lungospheres were of three different phenotypes: grape-like, cystic with a clearly defined lumen, and compound with regions resembling both the grape-like and the cystic phenotype. With FGF2-STAB, lungospheres were formed with the same phenotypes as with FGF2-wt, however, they formed with significantly higher LFE (0.132 ± 0.015%) (Figure1A, B; Figure S1B).

**Figure 1.**
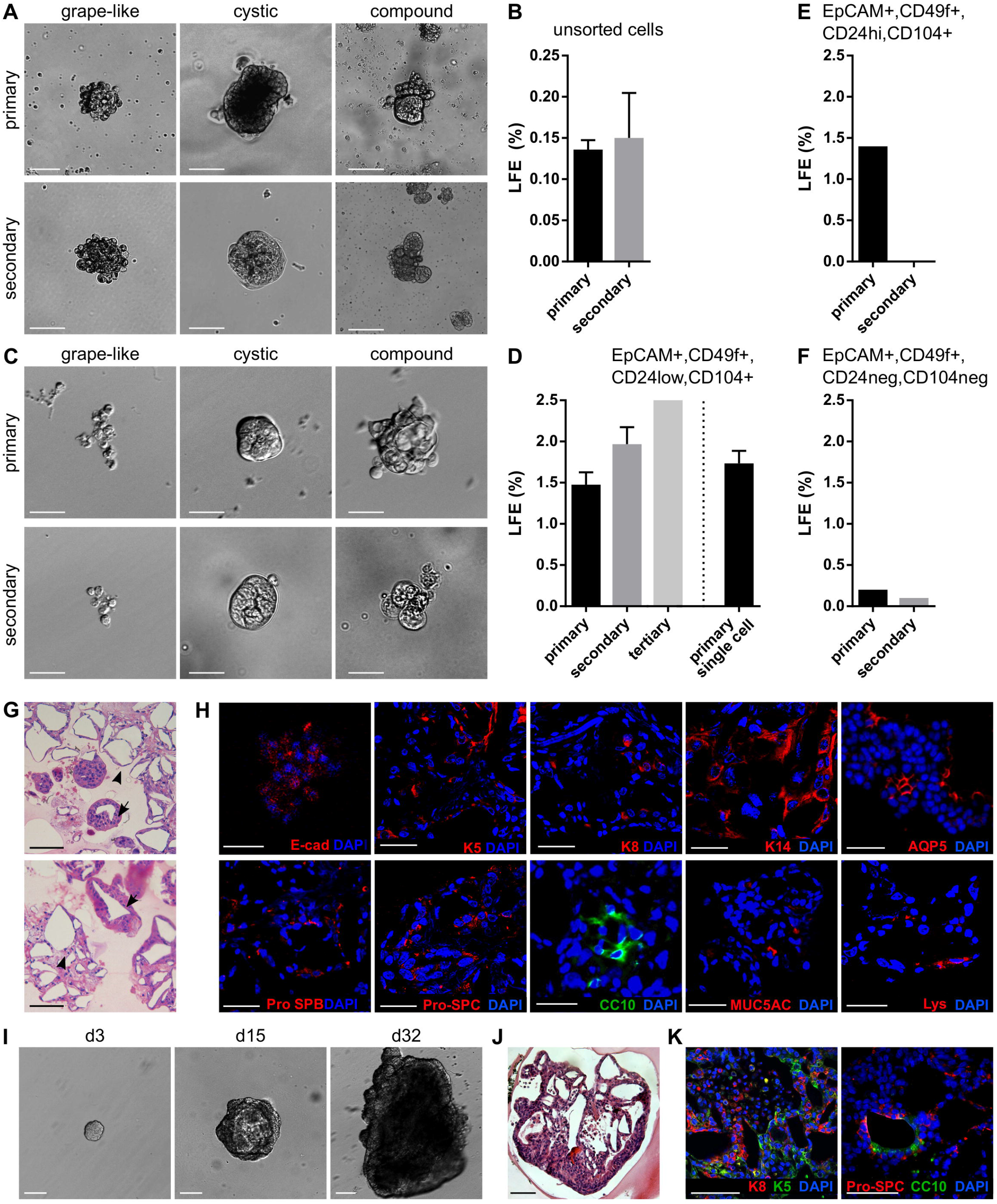
Lung epithelium contains LSPCs that form lungospheres with the capacity for self-renewal and differentiation. (**A, B**) Lungospheres formed from unsorted lung epithelial cells in nonadherent conditions with EGF and FGF2. (**A**) Representative photographs of primary and secondary lungospheres. Scale bars, 100 μm. (**B**) The efficiency of primary and secondary lungosphere formation with EGF and FGF2. The plots show mean + SD; n = 5. (**C, D**) Lungospheres formed from FACS-sorted EpCAM^+^,CD49f^+^,CD24^low^,CD104^+^ cells in non-adherent conditions with EGF and FGF2. (**C**) Representative photographs of primary and secondary lungospheres. Scale bar, 100 μm. (**D**) The efficiency of primary and secondary lungosphere formation with EGF and FGF2. The plots show mean + SD; n = 3 (n =1 for tertiary lungospheres). (**E, F**) The efficiency of primary and secondary lungosphere formation of FACS-sorted EpCAM^+^,CD49f^+^,CD24^hi^,CD104^+^ (**E**) and EpCAM^+^,CD49f^+^,CD24^neg^,CD104^neg^ (**F**) cells in non-adherent conditions with EGF and FGF2. The plots show mean + SD; n = 1. (**G, H**) Lungospheres grown in non-adherent conditions form lung-like structures. (**G**) The photographs of hematoxylin/eosin-stained paraffin sections show alveolar-like structures (arrowheads) and airways-like structures (arrows). (**H**) Immunofluorescence staining of paraffin sections for E-cadherin, keratin 5 (K5), keratin 8 (K8), keratin 14 (K14), Pro-SPB, Pro-SPC, aquaporin 5 (AQP5), CC10, MUC5AC, lysozyme (Lys), and nuclei (DAPI). (**I-K**) Lungospheres embedded in 3D Matrigel and cultured with EGF and FGF2 proliferate and form large organoids with lung-like structure. (**I**) The photographs show morphogenesis of a lungosphere-derived organoid over 32 days. (**J**) Hematoxylin/eosin-stained section of lungosphere-derived organoid. (**K**) Immunofluorescence staining of paraffin sections of lungosphere-derived organoids for keratin 5 (K5), cytokeratin 8 (K8), Pro-SPC, CC10, and nuclei (DAPI). (**G-K**) Scale bars, 100 μm.

To test the capacity of these spheres to self-renew, we performed a secondary lungosphere assay. Primary lungospheres were disintegrated to single cells and seeded again in non-adherent conditions. After 14 days of cultivation, secondary lungospheres of similar phenotypes as the primary lungospheres were formed at LFE 0.150 ± 0.055% (Figure1A, B), whereas the LFE in the presence of FGF2-wt was lower (0.100 ± 0.029%) (Figure S1C).

To investigate what cells of the lung epithelium are capable of lungosphere formation, we sorted the lung epithelial cells by FACS according to markers for adult lung stem cells (McQualter and Bertoncello, 2015) and cultured them in non-adherent conditions. The adult lung stem cell-enriched population characterized by EpCAM^+^,CD49f^+^,CD24^low^,CD104^+^ status (McQualter and Bertoncello, 2015) formed primary and secondary lungospheres of similar phenotypes as unsorted cells and with LFE 1.475 ± 0.150%) for primary and 1.967 ± 0.208% for secondary lungospheres, respectively (Figure 1C,D). In contrast, EpCAM^+^,CD49f^+^,CD24^hi^,CD104^+^ and EpCAM^+^,CD49f^+^,CD24^neg^,CD104^neg^ cells formed only primary lungospheres with LFE 1.4% and 0.2%, respectively, and failed to form secondary lungospheres (Figure 1E,F). These results indicated that the self-renewing lungosphere-forming stem cells are present in the EpCAM^+^,CD49f^+^,CD24^low^,CD104^+^ population.

To test whether the lungospheres were clonal and formed by division of single stem cells rather than being formed by cell aggregation, we sorted single EpCAM^+^,CD49f^+^,CD24^low^,CD104^+^ cells directly in 96-wells plates at only one cell per well in non-adherent conditions. We observed formation of lungospheres with LFE 1.733 ± 0.153% (Figure 1D), which was similar to the primary LFE when thousands of cells were cultured together in one well, suggesting that the lungospheres were clonal.

Histological analysis of lungospheres grown in non-adherent conditions revealed that lungospheres formed lung-like structures, with both alveolar-like morphologies and airways-like morphologies (Figure 1G). Using immunofluorescence analysis of the lungospheres we detected cells positive for lung epithelial cell markers cytokeratin 8 (K8) and E-cadherin, basal cell markers cytokeratin 5 (K5) and cytokeratin 14 (K14), club cell marker CC10, mucous cell marker MUC5AC, marker of secretory serous cells in the respiratory pathway lysozyme, markers of ATII cells prosurfactant protein B (pro-SPB) and prosurfactant protein C (pro-SPC), and the marker of the ATI cells aquaporin 5 (AQP5) (Figure 1H). Ultrastructural analysis by transmission electron microscopy showed that a lungosphere is composed of a group of cells connected by (Figure S2A). Some of the lungosphere cells have a dense cytoplasm that is rich in mitochondria and lamellar bodies (Figure S2A).

In summary, our observations suggest that lung epithelium contains LSPCs with the ability to resist anoikis, to self-renew and to form clonal spheres containing both stem/progenitor cells and more differentiated cells, and that the LSPCs can be efficiently isolated from lung using the lungosphere assay without the requirement for FACS.

### Lung epithelial cells form organoids in 3D Matrigel

Extracellular matrix (ECM) provides regulatory signals for epithelial tissue growth and patterning (De Arcangelis and Georges-Labouesse, 2000). Therefore, we embedded lungospheres formed in non-adherent conditions in 3D Matrigel and cultured them for over 30 days to investigate morphogenesis of lungospheres in physiologically more relevant setting. We observed formation of large cystic and/or branched organoids (Figure 1I). These organoids had a complex lung-like structure (Figure 1J). Ultrastructural analysis by transmission electron microscopy revealed different types of cells present in the lungosphere-derived organoids (Figure S2B). They included: ATI cell-like simple squamous cells with central nucleus and a thin cytoplasm, ATII cell-like cuboidal cells with central nucleus and cytoplasm rich in mitochondria and lamellar bodies, and club cell-like columnar cells with basally located nucleus and lamellar bodies within cytoplasm (Figure S2B). This cell diversity was further supported by the presence of variety of epithelial and lung cell markers, including K5, K8, Pro-SPC and CC10 (Figure 1K), as found by immunofluorescence.

Next, we investigated the ability of unsorted lung epithelial cells to form lung organoids directly in Matrigel from single cells. The cells were cultured in 3D Matrigel in the same medium as lungospheres. We observed formation of organoids with efficiency (organoid forming efficiency, OFE) of 0.110 ± 0.007% (Figure 2A,B), what was similar to the LFE of unsorted lung epithelial cells. Control experiments with FGF2-wt showed formation of lung organoids with similar phenotype but at lower OFE (0.067 ± 0.008%) (Supplementary Figure 3A,B).

**Figure 2.**
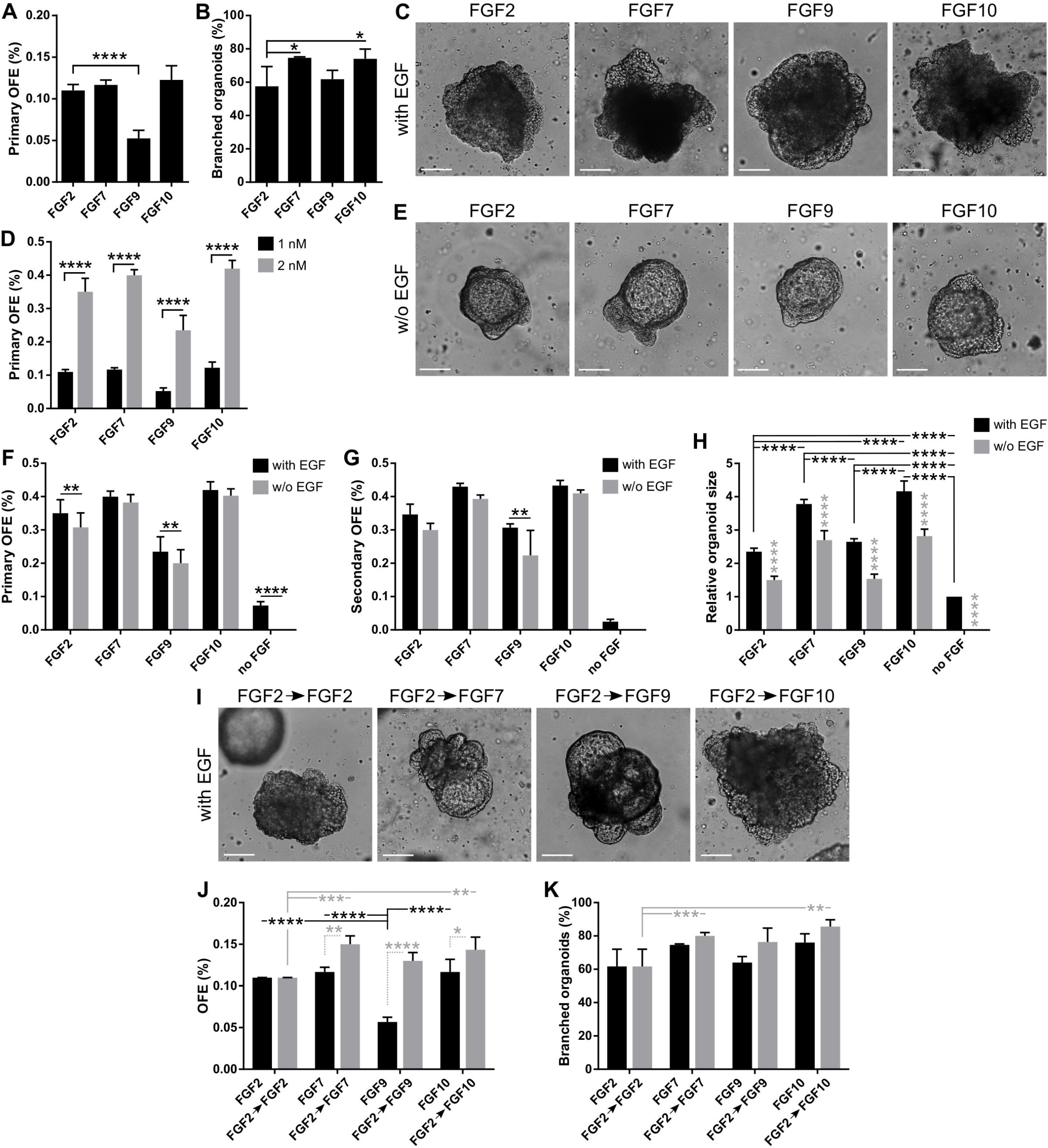
FGFs promote formation, proliferation and branching of lung organoids. (**A-C**) In 3D Matrigel, lung organoids are efficiently formed in response to different FGFs. (**A**) Primary lung organoid forming efficiency (OFE). The plot shows mean + SD; n = 3-5. ****P<0.0001 (one-way ANOVA). (**B**) Branching efficiency (%) of lung organoids, shown as mean + SD; n = 3-4. *P<0.05 (one-way ANOVA). (**C**) Representative photographs of primary lung organoids formed in 3D Matrigel in response to different FGFs (1 nM). Scale bars, 100 μm. (**D**) Increased concentration of FGFs increases organoid forming efficiency. The plot shows primary lung OFE as mean + SD; n = 3-5. ****P<0.0001 (two-way ANOVA). (**E-H**) EGF is not essential for lung organoid formation in the presence of FGFs. (**E**) Representative photographs of primary lung organoids formed in 3D Matrigel in the presence of different FGFs (2 nM) and in the absence of EGF. Scale bars, 100 μm. (**F, G**) The plots show primary (**F**) and secondary (**G**) lung organoid forming efficiency as mean + SD; n = 3-4. **P<0.01; ****P<0.0001 (two-way ANOVA). (**H**) Analysis of organoid size. The sizes are relative to organoids formed in medium with EGF and no FGF. The plots show mean + SD; n = 3, N = 15-20 organoids/treatment. The grey symbols indicate significance between the culture with and without EGF for the respective FGF. ****P<0.0001 (two-way ANOVA). (**I-K**) Initial treatment with FGF2 for 7 days increases lung organoid formation and branching efficiency in response to FGF7, FGF9 and FGF10. (**I**) Representative photographs of primary lung organoids formed in 3D Matrigel after 7 days of culture with 1 nM FGF2, followed by culture with different FGFs (1 nM). The pictures are from day 20 of culture. Scale bars, 100 μm. (**J, K**) The plots show primary lung OFE (**J**) and branching efficiency of lung organoids (**K**) as mean + SD; n = 3. *P<0.05; **P<0.01; ***P<0.001; ****P<0.0001 (two-way ANOVA).

### Different FGF ligands contribute differently to lung organoid formation

During lung development, different FGFs are differently required at specific stages of lung specification and morphogenesis (Volckaert and De Langhe, 2015). Therefore, we investigated the effect of different FGFs on lung organoid formation. To this end, we cultured unsorted lung epithelial cells in 3D Matrigel with EGF (20 ng/ml) and FGF2, FGF7, FGF9 or FGF10 (all at 1 nM). We found that with the different FGFs tested, the organoids formed with similar efficiencies as with FGF2 except for FGF9, with which the organoids formed with significantly lower efficiency (0.110 ± 0.007%, 0.117 ± 0.006%, 0.053 ± 0.017%, and 0.123 ± 0.017% for FGF2, FGF7, FGF9, and FGF10, respectively) (Figure 2A, C). Analysis of the organoid morphology revealed that FGF7 and FGF10, respectively, supported formation of more complex, branched structures in significantly higher proportion of organoids (Figure 2B). For all FGFs tested, increase of their concentration to 2 nM has led to at least twofold increase in numbers of produced organoids (Figure 2D).

Next, we tested the requirement of EGF for lung organoid formation. To this end, we cultured the lung epithelial cells in 3D Matrigel with media containing different FGFs (2 nM) in the presence (20 ng/ml) or absence of EGF. We found that lung organoids formed also in the absence of EGF (Figure 2E). However, removal of EGF from the cell culture media significantly reduced primary lung organoid formation in the case of culture with FGF2 and FGF9, and reduced secondary OFE in the case of FGF9 (Figure 2F, G). No organoids formed in the absence of both EGF and FGF. However, EGF on its own was capable of supporting lung organoid formation at low OFE (Figure 2F,G).

Analysis of the organoid morphology revealed that organoids that formed in the absence of EGF were significantly smaller compared to those formed with EGF (Figure 2H). Also, organoids formed with FGF7 and/or FGF10 were significantly bigger than the organoids formed with FGF2 and/or FGF9 (Figure 2H). Moreover, FGF7 and FGF10 induced organoid branching in significantly higher proportion of organoids than FGF2 or FGF9, irrespective of EGF presence (Supplementary Figure 4A,B). The absence of EGF significantly reduced branching of organoids formed with FGF9 (Supplementary Figure 4A). The most complex organoids with the highest number of branches were then produced by treatment with FGF10 (Supplementary Figure 4A,B).

We then investigated whether preculturing of lung epithelial cells with FGF2, followed by cultivation with different FGFs, would affect organoid formation and branching. We first cultured the lung epithelial cells for 7 days with FGF2 only and then continued with the culture in media containing FGF2, FGF7, FGF9, or FGF10 (Figure 2I). Overall, the preculture with FGF2 had positive effect on organoid formation. It rescued the inability of FGF9 to efficiently support organoid formation and also significantly increased organoid formation in medium with FGF7, FGF9, and FGF10, respectively (Figure 2I,J). In contrast to its effect on organoid formation, 7-day preculture with FGF2 did not influence organoid branching (Figure 2K). Still, the organoids cultured in FGF7 and FGF10, respectively, were superior in their branching capability compared to those cultured continuously in only FG2 (Figure 2K). This observation suggests that in the initial phase of lung organoid formation, cell survival and/or proliferation, which are efficiently supported by FGF2, might be the deciding factors for further organoid development. Then, in the later stage of lung organoid growth and patterning, organoid morphogenesis may be more driven by signals from FGF7, FGF9, and FGF10.

Collectively, our data indicate that at least *ex vivo*, FGF ligands assayed here may have partially redundant functions. An intriguing role seems to be played by FGF9 that has minimal capacity to promote organoid formation from single cells, but it is strong organoid-stimulating signal in later stages of their morphogenesis.

### Lung organoids contain both of alveolar-like and airway-like structures

FGF10 plays an important role during lung specification and morphogenesis (Ramasamy et al., 2007; Volckaert et al., 2013). In our experiments, FGF10 occurred as the most effective inducer of organoid formation and branching. Therefore, we focused our attention to the organoids formed with FGF10 for a closer characterization of their overall histological structure and cell type composition, as determined by hematoxylin-eosin staining and immunohistochemistry. Typically, we detected the presence of basally localized K5-positive cells and luminally localized K8-positive cells (Figure 3A,C). The organoids contained alveolar-like regions with cells expressing ATII and ATI cell markers pro-SPC and AQP5, respectively, as well as airway-like regions with ciliated cells (positive for acetylated tubulin) and club cells (positive for CC10) (Figure 3B).

**Figure 3.**
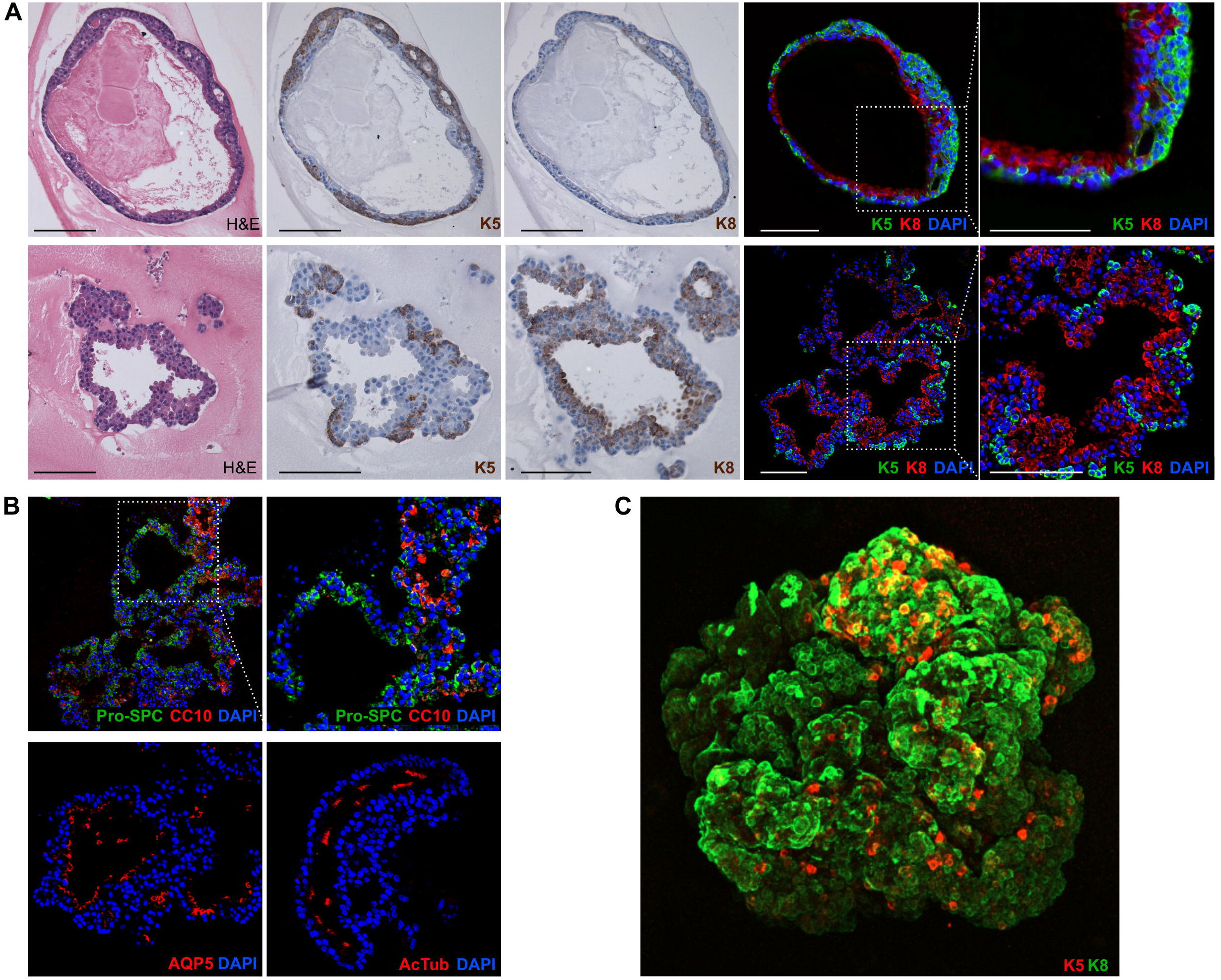
Lung organoids recapitulate cellular composition and organization of the lung. (**A**) Lung organoids have complex organization with basally localized keratin 5 positive cells and luminally localized keratin 8 positive cells. The photographs show paraffin sections of an organoid cultured in 3D Matrigel with FGF10, stained with hematoxylin/eosin (H&E) or by immunohistochemistry for keratin 5 (K5) or 8 (K8). Scale bars, 100 μm. (**B**) Lung organoids contain alveolar-like regions as well as airway-like regions. The photographs show immunofluorescence analysis of paraffin sections of lung organoids for markers of ATI cells (AQP5), ATII cells (Pro-SPC), ciliated cells (acetylated tubulin, AcTub), and club cells (CC10). Blue, nuclei (DAPI). Scale bars, 100 μm. (**C**) A whole-mount 3D confocal image of an organoid cultured in 3D Matrigel with FGF10, stained by immunofluorescence for keratin 5 (K5) and 8 (K8). Scale bars, 100 μm.

These results indicate that the lung organoid system, elaborated and used here, faithfully recapitulates the cellular composition and organization of the lung.

### WNT3A increases the efficiency of lung organoid formation and FGF10 promotes differentiation to lung epithelial cell lineages

WNT signaling plays an essential role during lung development (Frank et al., 2016) and repair after injury (Whyte et al., 2012; Lee et al., 2017). Moreover, WNT signaling is critically important for self-renewal and specification of stem cells in multiple organs (Clevers et al., 2014) and has been successfully employed in a protocols for production of organoids from several organs, including lungs (Heo et al., 2018). Therefore, here we also investigated the formation of lung organoids under the conditions involving WNT signals.

The WNT-based protocols typically combine WNT3A, R-Spondin1 (WNT agonist), Noggin (bone morphogenic protein inhibitor), and EGF, and optionally also FGFs, to support organoid growth in basement membrane ECM. WNT3A can be supplemented to the medium in the form of recombinant protein from a commercial supplier, or in the form of WNT3A-conditioned medium (WCM; (Sugimoto and Sato, 2017). We tested both approaches.

WNT3A efficiently promoted lung organoid formation but WCM was significantly less effective in this phenomenon (Figure 4A,B). Moreover, in the culture variant with WCM, we observed a striking new phenotype – colonies of cells of mesenchymal-like morphology (Figure 4A,C). These colonies started off as epithelial structures but changed into mesenchymal-like cells, intriguingly resembling the process of epithelial-to-mesenchymal transition (EMT). We hypothesized that factor(s) responsible for such effect of WCM may originate from the serum contained in WCM preparation as part of the culture medium. Therefore, we prepared also serum-free WCM (WCM-I) by culturing cells in medium containing ITS (insulin-transferrin-selenium) instead of serum. As expected, the EMT-like phenotype was much less pronounced in the WCM-I compared to serum-containing WCM (WCM-S). Also, WCM-I was superior to WCM-S in the efficiency of organoid production (Figure 4A-C).

**Figure 4.**
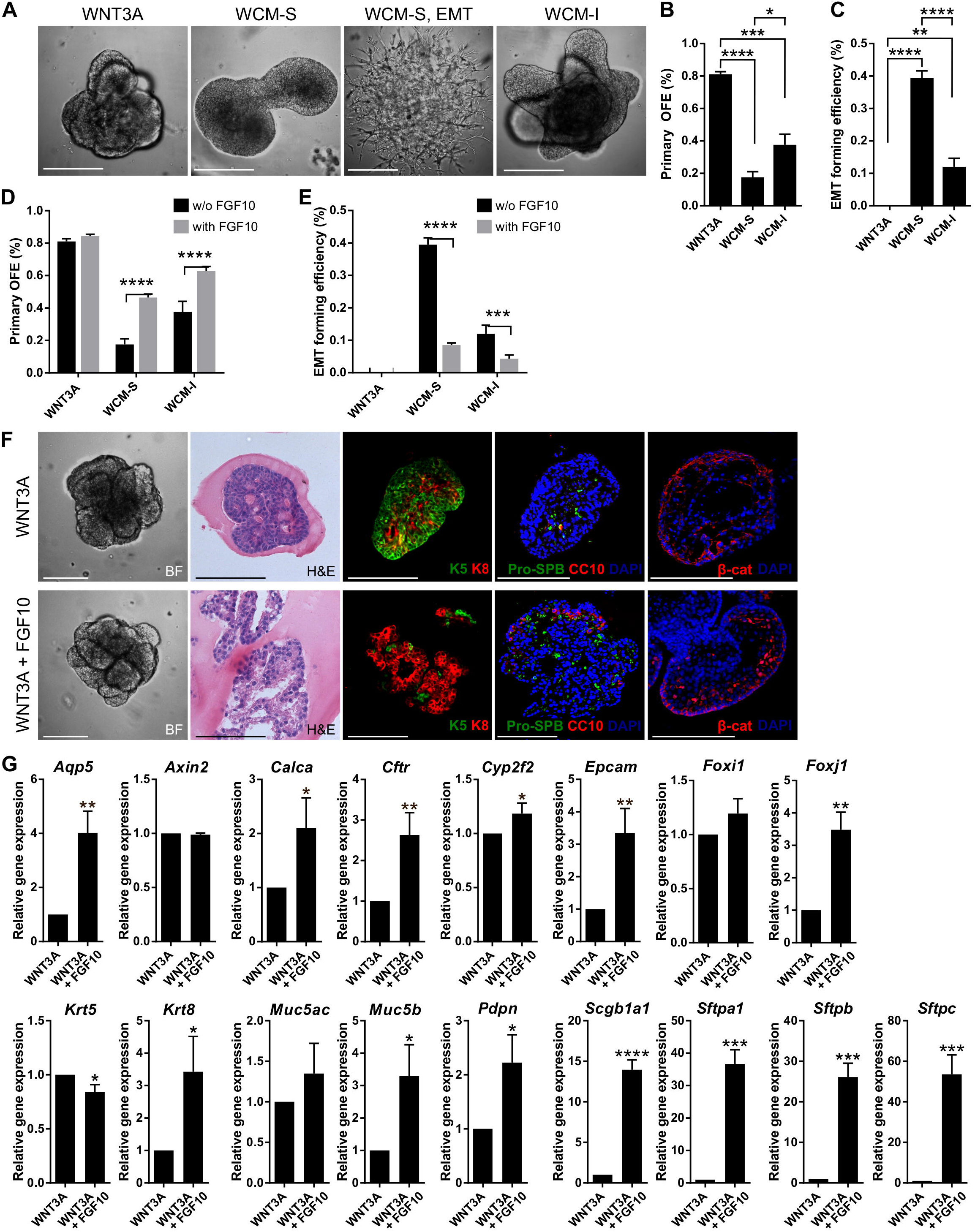
WNT3A enhances the efficiency of lung organoid formation and addition of FGF10 promotes organoid differentiation. (**A**) Representative photographs of lung organoids formed with WNT-based protocol in 3D Matrigel. WCM-S, WNT3A-conditioned medium with serum; WCM-I, serum-free WNT3A-conditioned medium (with ITS); EMT, epithelial to mesenchymal transition. (**B, C**) Primary lung organoid forming efficiency (OFE; **B**) and efficiency of formation of EMT colonies (**C**). The plots show mean + SD; n = 3. *P<0.05; **P<0.01; ***P<0.001; ****P<0.0001 (one-way ANOVA). (**D, E**) FGF10 increases lung organoid formation efficiency and decreases EMT occurrence in the cultures with WCM-based media. The plots show mean + SD; n = 3. ***P<0.001; ***P<0.001 (two-way ANOVA). (**F**) Brightfield images (BF), histological analysis (H&E) and immunofluorescence staining of lung organoids. Scale bar, 100 μm. (**G**) Results from qPCR analysis of expression of candidate genes of WNT3A treatment with or without FGF10. The plots show mean + SD, n = 3. *P<0.05; **P<0.01; ***P<0.001; ****P<0.0001 (Student’s t-test).

Because of the important role of FGF10 during lung epithelial development, demonstrated also here, we further investigated the effect of FGF10 on formation of lung organoids under conditions utilizing WNT signaling. Curiously, while FGF10 did not potentiate organoid forming efficiency when combined with WNT3A, it did significantly increase organoid formation in cultures containing either type of WCM (Figure 4D). FGF10 also significantly reduced occurrence of EMT-like phenotype in WCM-containing cultures (Figure 4E).

Histological and immunohistochemical analysis of lung organoids produced under conditions containing WNT3A alone or WNT3A plus FGF10, respectively, demonstrated that the presence of FGF10 dramatically increased a degree of organoid differentiation. Specifically, organoids grown under the influence of FGF10 contained expanded lung-like regions (Figure 4F) and typically expressed much higher numbers of Pro-SPC- and CC10-positive cells (Figure 4F). The organoids formed with WNT3A only were more compact and contained several layers of less differentiated basal cells (Figure 4F). It is of note that organoids produced under conditions containing WNT3A showed nuclear localization of β-catenin, thus documenting activation of WNT signaling pathway (Figure 4F).

We have further approached the regulatory significance of WNT and FGF10 by quantifying the expression of 16 genes that define specific types of cells and/or stages of development of lung epithelial cell lineages (Figure 4G). To this end, the organoids were produced in the media containing WNT3A with or without FGF10. The key findings were as follows. The organoids expressed all the genes assayed, however, the expression levels were influenced by the presence and absence of FGF10. Although the statistical significance of the differences varied among the individual genes, overall, the expression of genes associated with differentiated cell phenotypes was higher in organoids formed in FGF10-containing medium. Specifically, this difference was very highly pronounced for AQP5 (*Aqp5*; ATI cells), CFTR (*Cftr*; anion secretory cells), EpCAM (*Epcam*; epithelial cells), FoxJ1 (*Foxj1*; ciliated cells), CC10 (*Scgb1a1*; club cells), SP-A (*Sftpa1*; ATII cells), SP-B (*Sftpb*; ATII cells) and SP-C (*Sftpc*; ATII cells), less pronounced for CGRP (*Calca*; neuroendocrine cells), P450 (*Cyp2f2*; club cells), CFTR (Cftr; anion secretory cells), K8 (*Krt8*; epithelial cells), MUC5B (*Muc5b*; mucosa cells) and PDPN (*Pdpn*; ATI cells), and it was statistically insignificant for FOXI1 (*Foxi1*; pulmonary ionocytes) and MUC5AC (*Muc5ac*; mucosa cells). Importantly, FGF10 had opposite effect on K5 (*Krt5*; basal cells), with this stemness-associated gene being more expressed in organoids produced in the absence of FGF10. It is of note that FGF10 did not influence intracellular action of WNT3A, as indicated by sustained expression of WNT target *Axin2* (Figure 4G).

Together, this set of data identify WNT3A as a factor that preferentially enhances organoid formation, whereas FGF10 drives differentiation of cells within the organoids.

## Discussion

FGF and WNT signaling pathways are essential components of the gene regulatory network in the lungs. They coordinate maintenance of stem/progenitor cells, epithelial/mesenchymal patterning, and branching morphogenesis during lung development and repair (Volckaert and De Langhe, 2015). In this study, we investigated the roles of FGF ligands and also WNT3A in lung epithelial morphogenesis using 3D cell culture model that we have developed.

Our study revealed that unsorted lung epithelial cells as well as sorted LSPCs (EpCAM^+^ CD49f^+^ CD24^low^ CD104^+^) are capable of proliferation, self-renewal and differentiation in non-adherent conditions to form spheres, similar to epithelial stem cells from other organs (Reynolds and Weiss, 1992; Shaw et al., 2012; Zhao et al., 2015). The lungospheres formed structures that could be categorized into three phenotypes, which were similar to the phenotypes described for LSPC-derived organoids grown in ECM (McQualter et al., 2010), suggesting that the mechanisms underlying these phenotypes are independent of the ECM. However, embedding of the lungospheres in 3D ECM (Matrigel) supported further development, including growth and morphological changes of the lungospheres into lung organoids with branched lung-like structures.

Direct seeding of unsorted lung epithelial cells into Matrigel promoted organoid formation at similar efficiency as culture in non-adherent conditions, suggesting that the lungospheres and lung organoids come from the same type of cells. Importantly, while several studies reported that co-culture with stromal cells is required for distal LSPCs to form lung-like epithelial structures (McQualter et al., 2010; Chen et al., 2012; Barkauskas et al., 2013; Lee et al., 2014; Hegab et al., 2015), our cell culture method in defined medium with EGF and FGF ligands enables lung organoid formation without the need for stromal cells. Several studies reported the ability of FGF ligands to replace the requirement for stromal support (McQualter et al., 2010; Hegab et al., 2015). Interestingly, in these studies, 5 to 10 times higher concentration of FGF was used than in our study, suggesting that our cell culture protocol is more effective. Our study also shows the advantage of using hyperstable growth factor variants, such as FGF2-STAB, for lung organoid culture, and extends the cell culture potential of FGF2-STAB beyond the reported use in human embryonic stem cell culture (Dvorak et al., 2018).

Our experiments addressing organoid forming efficiency and organoid morphology revealed that FGF ligands FGF2, FGF7, FGF9, and FGF10 act partially redundantly *ex vivo*. All tested FGF ligands were capable of supporting organoid formation and branching, with FGF9 showing a significantly lower capacity to promote lung organoid formation compared to other FGF ligands. However, preculture with FGF2 followed by further culture with FGF9 was able to rescue the low organoid formation, typical for treatment with only FGF. Moreover, preculture with FGF2 further increased the organoid forming efficiency of FGF7, FGF9, and FGF10, suggesting that FGF2 supported cell survival and/or proliferation more efficiently than other FGF ligands. In contrast to our *ex vivo* observations, only FGF10 is essential for lung development *in vivo*. FGF10 null mice show complete lung agenesis (Min et al., 1998; Sekine et al., 1999), whereas inactivation of FGF2, FGF7 or FGF9 affects only lung patterning and branching morphogenesis (Zhou et al., 1998; Guo et al., 1996; White et al., 2006).

WNT signaling plays important roles in self-renewal and differentiation of stem cell in adults as well as during embryonic development (Mucenski et al., 2003; Clevers et al., 2014; Bhavanasi and Klein, 2016). WNT3A was reported to potentiate alveolar organoid formation from ATII cells and also expansion of ATII cells *ex vivo* (Frank et al., 2016; Lee et al., 2017). Furthermore, WNT activation promotes clonal expansion of ATII cells during alveologenesis *in vivo* (Frank et al., 2016). In this study, we showed that WNT3A significantly increases organoid forming efficiency of unsorted lung epithelial cells in comparison to WNT3A-free conditions. Importantly, we report on major differences in organoid forming efficiency depending on the source of WNT3A. Recombinant WNT3A demonstrated the highest capacity to promote organoid formation, while conditioned media from cells expressing WNT3A showed lower capacity to promote organoid formation. While the lower capacity to support organoid formation could, in theory, be compensated by increasing the proportion of WNT3A-conditioned medium in the organoid medium, it would not bring much advantage. And that is because the WNT3A-conditioned medium also induces formation of disorganized colonies of mesenchymal-like cells. These colonies arise probably through epithelial-to-mesenchymal transition (EMT) of lung epithelial cells because we frequently observed transition of smooth, epithelial-like organoid structures to mesenchymal-like colonies of spiky morphology. The L-cell-derived factors that induce such EMT-like behavior remain to be determined.

When FGF10 was added to the WCM-based medium, it increased the efficiency of organoid formation and decreased the incidence of mesenchymal-like structures. However, when FGF10 was added to the medium with recombinant WNT3A, it did not increase the formation of organoids any further, suggesting that WNT3A saturated organoid formation to its maximum. Still, we observed significantly higher expression of markers of all lung cell lineages, suggesting that FGF10 promoted organoid differentiation to both airway and distal lung epithelium. This is consistent with the reported role of FGF10 in regulation of lung progenitor differentiation based on the developmental context (Volckaert et al., 2013) as well as during homeostasis and in regeneration (Volckaert et al., 2017; Yuan et al., 2019).

3D culture models of the lung, including lung organoids represent an invaluable tool for developmental biology, cancer biology, pharmacology and disease modelling (Barkauskas et al., 2017; Rabata et al., 2017). They offer wide range of modalities and can be adjusted according to the research needs. In this work, we developed and utilized lung organoid model based on unsorted lung epithelial cells cultured in Matrigel in defined medium to assess the role of FGF and WNT signaling in lung epithelial morphogenesis. Our findings contribute to better understanding of the complex signaling pathways involved in lung organogenesis with potential applications in tissue engineering and regenerative medicine. Furthermore, as demonstrated, our model can be used for testing of growth factor variants created by protein engineering and potentially other proteins or bioactive substances, providing that the targeted pathways are involved in lung epithelial morphogenesis, including cell survival, proliferation or differentiation.

## Materials and Methods

### Mice

Female ICR mice of 6-10 weeks of age were used as donors of the lung tissue. The mice were housed, handled and sacrificed in accordance with the Czech and European laws for the use of animals in research, under a valid project license at the Laboratory animal breeding and experimental facility of Masaryk University.

### Isolation of lung epithelial cells by differential centrifugation

Lung epithelial cells were isolated as described previously (Rabata et al., 2017). Briefly, the lungs were mechanically disintegrated using scalpels and digested by a solution of collagenase and trypsin [2 mg/ml collagenase A, 2 mg/ml trypsin, 5 μg/ml insulin, 50 μg/ml gentamicin (all Sigma/Merck), 5% fetal bovine serum (FBS; Hyclone/GE Healthcare) in DMEM/F12 (Thermo Fisher Scientific)] for 45 min at 37°C with shaking. The resulting tissue suspension was centrifuged and the pellet was treated with Red blood cells lysis buffer [155 mM NH4Cl (Penta), 12 mM NaHCO3 (Fluka), 0.1 mM EDTA (Merck) in distilled H2O, pH 7.4, filter sterilized]. After washing with DMEM/F12 and centrifugation, the cell pellet was treated by DNase I (20 U/ml; Merck). After final wash with DMEM/F12, the cell pellet was exposed to five rounds of differential centrifugation to remove mesenchymal cells. The resulting pellet of primary lung epithelial organoids was further processed by incubation in HyQTase (Hyclone/GE Healthcare) and repetitive pipetting to produce single-celled suspension.

### Isolation of lung epithelial stem/progenitor cells by FACS

LSPCs were isolated as described previously (McQualter and Bertoncello, 2015; Rabata et al., 2017). Briefly, the lungs were chopped up using scalpels and the resulting mince was digested by Liberase (0.048mg/ml; Roche) and incubated for 45 min at 37°C with shaking. The suspension was passed through 18G and 21G needles and treated with DNase I (20 U/ml). Next, the suspension was passed through a 100 μm cell strainer, treated with Red blood cell lysis buffer and passed through a 40 μm cell strainer. The resulting single-celled suspension was centrifuged to collect the cells. The pellet was suspended in blocking buffer [1% BSA in HBSS (both Merck)] and incubated for 20 min at room temperature. After centrifugation, the cells were resuspended at 1 × 10^7^ cells/ml in FACS buffer containing the selection antibody cocktail (anti-CD104, anti-EpCAM, anti-CD24, anti-CD49f, anti-CD45; see Supplementary Table 1) and incubated in ice in the dark for 20 min. The cells were washed with HBSS, passed through a 30 μm cell strainer and incubated with 10 μl/ml 7-ADD (BD Biosciences) for 5 min in ice in the dark. Then cells were sorted using FACSAria II SORP (BD Biosciences).

### Lungosphere culture in non-adherent conditions

Lung epithelial cells were cultured in non-adherent conditions as described previously (Rabata et al., 2017). Briefly, for primary lungosphere assay, unsorted lung epithelial cells were seeded in polyHEMA-treated 6-well plates at 2.5 × 10^4^ to 5 × 10^4^ cells in 2 ml/well of lungosphere medium [1 × B-27 without vitamin A, 100 U/ml penicillin, 100 μg/ml streptomycin (all Thermo Fisher Scientific), 4 μg/ml heparin (Sigma/Merck), 20 ng/ml EGF (Peprotech), 10 ng/ml FGF2-STAB (thermostable FGF2; Enantis) or 10 ng/ml FGF2-wt (Peprotech), 10 μM Y-27632 (Merck) in phenol red-free DMEM/F12 (Thermo Fisher Scientific)], or 500 to 1000 FACS-sorted cells were seeded in polyHEMA-treated 24-well plates in 1 ml/well of lungosphere medium. The plates were incubated in a humidified atmosphere at 37°C, 5% CO_2_. Fresh lungosphere medium (without Y-27632) was added every 3 days. Lungospheres were counted after 10 to 15 days of culture. To passage lungospheres (for secondary and tertiary lungosphere assay), the lungospheres were collected from the plates and processed to single-celled suspension by HyQTase digestion and repetitive pipetting. Resulting single cells were reseeded in polyHEMA-treated plates in lungosphere medium at the same density as for primary assay. Lungosphere forming efficiency (LFE, %) was calculated as (number of lungospheres formed)/(number of cells seeded) × 100.

### Embedded culture of lungospheres in 3D Matrigel

The lungospheres were collected from the polyHEMA-treated plates, washed with DMEM/F12 and then with basal culture medium [1× ITS (10 μg/ml insulin, 5.5 μg/ml transferrin, 6.7 ng/ml selenium), 100 U/ml penicillin, 100 μg/ml streptomycin in DMEM/F12 (all Thermo Fisher Scientific)]. Then the lungospheres were mixed with Matrigel (growth factor reduced; Corning) and plated into Matrigel-coated 24-well plate in domes. The plate was incubated at 37°C for 30-45 min before adding basal culture medium supplemented with growth factors as needed. The plate was incubated in a humidified atmosphere of cell culture incubator (37°C, 5% CO_2_). Medium was changed every 2-3 days.

### Lung organoid culture in 3D Matrigel

The unsorted or sorted lung epithelial cells were resuspended in Matrigel and plated into Matrigel-coated 24-well plate in domes (1-2 × 10^4^ cells in 50 μl Matrigel/well). The plate was incubated at 37°C for 30-45 min before adding 1 ml of medium per well. The cells were incubated in a humidified atmosphere of cell culture incubator (37°C, 5% CO_2_). Medium was changed every 3 days. The media used were: Lungosphere medium: [1× B-27 without vitamin A, 100 U/ml penicillin, 100 μg/ml streptomycin, in phenol red-free DMEM/F12], supplemented with EGF (20 ng/ml) and/or FGFs [1 or 2 nM FGF2-STAB, FGF2-wt, FGF7, FGF9, or FGF10] as needed according to experiment, and 10 μM Y-27632 (only for the first 3 days of culture). WNT lung organoid medium (Lee et al., 2017): 50 ng/ml WNT3A (Peprotech) or 50% WNT3A-conditioned medium, 100 ng/ml mouse Noggin, 500 ng/ml human R-spondin 1 (both Peprotech) and 40 ng/ml EGF in lungosphere medium, with or without 40 ng/ml FGF10. Organoid forming efficiency (OFE, %) was calculated as (number of organoids formed)/(number of cells seeded) × 100.

### Production of WNT3A-conditioned medium

WNT3A-conditioned medium (WCM) was prepared using the cell line L WNT3A (ATCC®, CRL-2647™) (Sugimoto and Sato, 2017). The L cells were cultured in DMEM medium (Thermo Fisher Scientific), 100 U/mL penicillin, 100 μg/mL streptomycin with 10% FBS or 1× ITS. The medium was collected and sterile filtered after three days of culture (first batch), then fresh medium was added to the cells for another two days until the medium was collected and sterile filtered (second batch). The first and second batches of medium were mixed (1:1), resulting in the WCM. The WCM was aliquoted and stored at −20°C until used.

### Histological and immunohistochemical analysis

The lungospheres formed in non-adherent conditions or organoids formed in 3D Matrigel were fixed with 4% paraformaldehyde in PBS for 30 min, washed with PBS and embedded in 3% low melting point agarose (Merck). Then the samples were processed via standard procedure for paraffin embedding. Paraffin sections were cut (2 μm thickness), deparaffinized using xylene and rehydrated. For histological analysis, the sections were stained with hematoxylin and eosin, dehydrated and mounted in Pertex (Histolab Products). For immunohistochemistry analysis, antigens were retrieved using Citrate buffer (Dako) for 30 min and endogenous peroxidase activity was blocked using 3% hydrogen peroxide. The sections were blocked in PBS with 10% FBS and incubated with primary antibody (Supplementary Table 1) for 1 h at RT. After washing, sections were incubated with secondary antibody (anti-mouse, EnVision+ Dual Link System-HRP; Dako) for 30 min at RT. After washing, bound secondary antibody was detected using Liquid DAB+ Substrate Chromogen System (Dako). The nuclei were stained with Mayer’s hematoxylin, dehydrated and mounted in Pertex. The photographs were taken using Leica DM5000B microscope equipped with Leica DFC480 camera. For immunofluorescence analysis, the sections were immersed in Citrate buffer and blocked with PBS with 10% FBS. Then the sections were incubated with primary antibodies (Supplementary Table 1) overnight at 4°C. After washing, the sections were incubated with secondary antibodies (Supplementary Table 1) for 2 h at RT. Then sections were washed, stained with DAPI (1 μg/ml; Merck) for 10 min and mounted in Mowiol (Merck). Fluorescence was detected and documented using Nikon Eclipse Ti2 inverted microscope or using Olympus FV500 and FV3000 confocal laser scanning microscope.

### Real-time quantitative PCR (qPCR)

RNA was isolated using RNeasy Mini Kit (Qiagen) according to the manufacturer’s instruction. cDNA was prepared using High Capacity RNA-to-cDNA kit (Thermo Fisher Scientific). Real-time qPCR was performed using 5 ng cDNA, 5 pmol of the forward and reverse gene-specific primers each (primer sequences shown in table S2) in Light Cycler SYBR Green I Master mix (Roche) on LightCycler 480 II (Roche). Relative gene expression was calculated using the ΔΔCt method and normalization to two housekeeping genes, β-actin (*Actb*) and Eukaryotic elongation factor 1 γ (*Eef1g*).

### Statistics

Statistical analysis was performed using the Prism software (GraphPad) using Student’s t-test and ANOVA. *P < 0.05, **P < 0.01, ***P < 0.001, ****P < 0.0001. The number of independent biological replicates is indicated as *n*.

## Ethics Statement

The animal study was reviewed and approved by the Ministry of Agriculture of the Czech Republic, and the Expert Committee for Laboratory Animal Welfare at the Faculty of Medicine, Masaryk University.

## Author Contributions

A.R. performed the experiments, analyzed the data and drafted the manuscript. Z.K. conceptualized the study, designed the experiments, analyzed the data and wrote the manuscript. R.F. and K.S. performed sorting of cells by FACS. A.H. secured funding and revised the manuscript. All authors approved the final manuscript.

## Conflict of Interest Statement

The authors declare that the research was conducted in the absence of any commercial or financial relationships that could be construed as a potential conflict of interest.

## Abbreviations

AQP5: aquaporin 5
ATI: alveolar type I cell
ATII: alveolar type I cell
CC10: club cells specific 10 protein
ECM: extracellular matrix
EGF: epithelial growth factor
EMT: epithelial to mesenchymal transition
FACS: fluorescence-activated cell sorting
FGF: fibroblast growth factor
FGF2-STAB: stabilized FGF2
FGF2-wt: wild-type FGF2
FGFR: fibroblast growth factor receptor
K5: keratin 5
K8: keratin 8
LFE: lungosphere forming efficiency
LSPC: lung epithelial stem/progenitor cells
OFE: organoid forming efficiency
pro-SPB: pro-surfactant protein B
pro-SPC: pro-surfactant protein C
WCM: WNT3A-conditioned medium.

## Acknowledgements

We thank Prof. Vitezslav Bryja for providing WNT3A-producing L cells, Dr. Jana Dumkova for help with electron microscopy, and Katarina Mareckova for excellent histology service. We acknowledge the core facility CELLIM of CEITEC supported by the Czech-BioImaging large RI project (LM2015062 funded by MEYS CR) for their support with obtaining scientific data presented in this paper.

## Funding

This study was supported by funds from the Ministry of Health of the Czech Republic (Grant 16-31501A to A.H.), the Ministry of Education, Youth, and Sports of the Czech Republic (National Program of Sustainability II, project no. LQ1605 to A.H. and K.S.), and from the Faculty of Medicine of Masaryk University (project no. MUNI/A/1382/2019).

## Contribution to the Field Statement

Respiratory diseases and cancers are among the top ten leading causes of death around the world. For development of new treatments and regenerative strategies, understanding of the regulation of lung epithelial morphogenesis and homeostasis is essential. In this work, we investigated the roles of FGF and WNT signaling in formation of lung stem cell-derived organoids and their morphogenesis using 3D culture models, immunostaining and imaging. We reveal previously unreported partially redundant roles of FGF ligands in lung organoid formation and branching, and the roles of FGF10 and WNT signaling in balancing stem cell expansion and differentiation *ex vivo*. Moreover, we report important application notes towards the use of hyperstable growth factors and recombinant proteins versus conditioned media. Our findings provide stepping stones towards development of robust lung organoid models for basic research, disease modeling and drug development, and towards lung tissue engineering.

