## Supplementary Material for "3D cell culture models demonstrate a role for FGF and WNT signaling in regulation of lung epithelial cell fate and morphogenesis"

### Supplementary Methods

#### Transmission electron microscopy

The lungospheres that formed in non-adherent conditions or in 3D Matrigel were washed in 0.1 M cacodylate buffer [0.1 M sodium cacodylate in distilled water (Sigma/Merck)], fixed with 3% glutaraldehyde for 1 h and postfixed in 1% $OsO_4$ for 50 min. The cells were washed in cacodylate buffer, embedded in 1% agar blocks, dehydrated in increasing series of ethanol (50, 70, 96, and 100%), treated with 100% acetone, and embedded in Durcupan resin (Sigma/Merck). Ultrathin sections were prepared using LKB 8802A Ultramicrotome, stained with uranyl acetate and Reynold's lead citrate (Sigma/Merck), and examined with FEI Morgagni 286(D) transmission electron microscope.

### Supplementary Figures and Tables

### Supplementary Figures

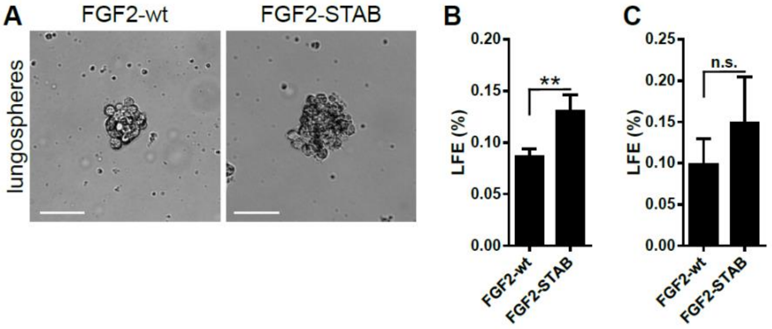


**Supplementary Figure 1. Hyperstable FGF2 promotes formation of lungospheres with higher efficiency than wild-type FGF2.** (**A**) Representative photographs of lungospheres formed in the presence of FGF2-wt and FGF2-STAB. Scale bars, 100 µm. (**B, C**) The efficiency of primary (**B**) and secondary (**C**) lungosphere formation (LFE). The plots show mean + SD; n = 4-6 (Student's t-test).

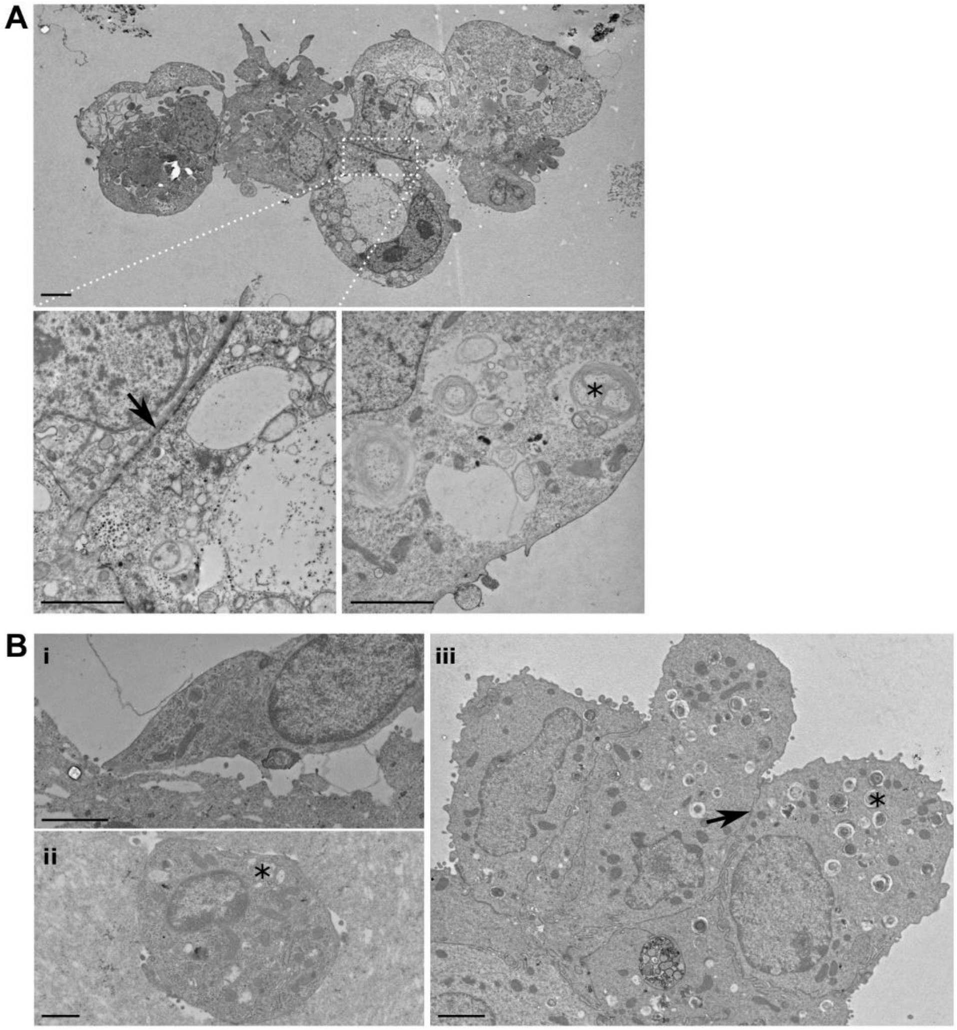


**Supplementary Figure 2. Ultrastructural analysis of lungospheres and lung organoids**. (**A**) Transmission electron microscopy (TEM) photograph of a lungosphere formed from unsorted lung epithelial cells in non-adherent conditions. The cells attach to each other by intercellular junctions (indicated by an arrow). Some cells contain lamellar bodies (indicated by a star). (**B**) TEM photographs of cells from lung organoids cultured in 3D Matrigel. The photographs show (i) ATI-like cell, (ii) ATII-like cell, and (iii) club cell-like cell. Arrow, intercellular junctions; stars, lamellar bodies. Scale bars, 2 μm.

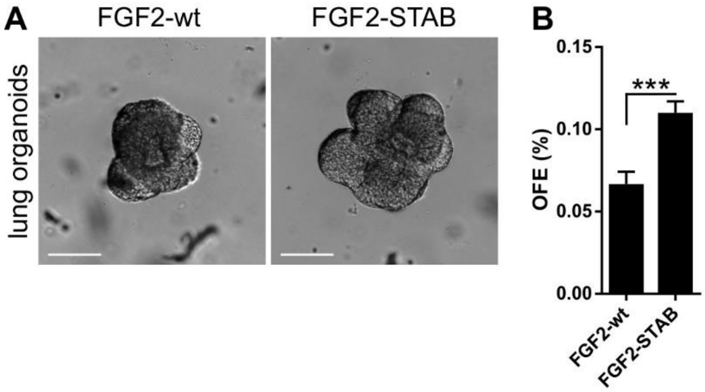


**Supplementary Figure 3. Hyperstable FGF2 promotes formation of lung organoids with higher efficiency than wild-type FGF2.** (**A**) Representative photographs of lung organoids formed in the presence of FGF2-wt and FGF2-STAB. Scale bars, 100 μm. (**B**) The efficiency of primary lung organoid formation (OFE), shown as mean + SD; n = 3 (Student's t-test).

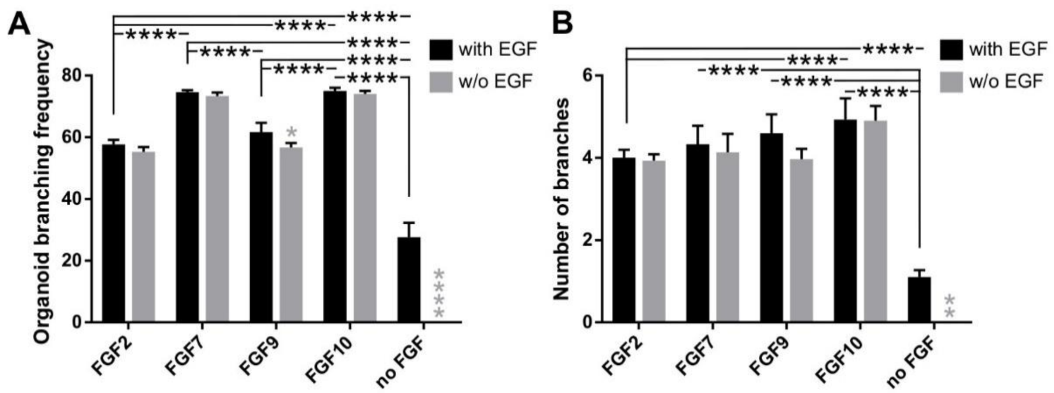


**Supplementary Figure 4.** (**A, B**) **EGF is not essential for lung organoid formation in the presence of FGFs.** The plots show organoid branching frequency (**A**) and number of branches (**B**) of organoids formed in media with different FGFs and with or without EGF as indicated. The plots show mean + SD; n = 3, N = 15-20 organoids/treatment. The grey symbols indicate significance between the culture with and without EGF for the respective FGF. *$P<0.05$; **$P<0.01$; ****$P<0.0001$ (two-way ANOVA).

### Supplementary Tables

**Supplementary Table 1.** Overview of antibodies used in this study.

| Antigen | Clone | Catalog # | Conjugate | Host | Supplier | Dilution |
|---|---|---|---|---|---|---|
| **Flow cytometry:** | | | | | | |
| CD24 | M1/69 monoclonal | 48-0242-82 | eFluor 450 | Rat | eBioscience | 1:1000 |
| CD45 | 30-F11 monoclonal | 25-0451-82 | PECy7 | Rat | eBioscience | 1:1000 |
| CD49f | GoH3 monoclonal | 12-0495-82 | PE | Rat | eBioscience | 1:1000 |
| CD104 | 346-11A monoclonal | 123608 | Aleax Fluor 647 | Rat | BioLegend | 1:1000 |
| EpCAM | G8.8 monoclonal | 11-5791-82 | FITC | Rat | eBioscience | 1:1000 |
| 7-AAD | N/A | 559925 | IP | N/A | BD Biosciences | 1:100 |
| **Immunofluorescence:** | | | | | | |
| Keratin 5 | Poly19055 polyclonal | 905504 | Unconjugated | Rabbit | BioLegend | 1:200 |
| Cytokeratin 8 | 1E8 | 904804 | Unconjugated | Mouse | BioLegend | 1:200 |
| Keratin 14 | Poly19053 polyclonal | 905304 | Unconjugated | Rabbit | BioLegend | 1:200 |
| E-cadherin | M168 monoclonal | ab76055 | Unconjugated | Mouse | Abcam | 1:200 |
| Prosurfactant protein B | Polyclonal | ab15011 | Unconjugated | Rabbit | Abcam | 1:200 |
| Prosurfactant protein C | Polyclonal | AB3786 | Unconjugated | Rabbit | Millipore | 1:200 |
| CC10 | T-18 polyclonal | sc-9772 | Unconjugated | Goat | Santa Cruz | 1:200 |
| Aquaporin 5 | G-19 polyclonal | sc-9890 | Unconjugated | Goat | Santa Cruz | 1:100 |
| MUC5AC | 45M1 monoclonal | MA5-12178 | Unconjugated | Mouse | Thermo Fisher Scientific | 1:100 |
| Aceylated α tubulin | 6-11B-1 monoclonal | sc-23950 | Unconjugated | Mouse | Santa Cruz | 1:100 |
| Lysozyme | EPR2994(2) monoclonal | ab108508 | Unconjugated | Rabbit | Abcam | 1:200 |
| β-catenin | 12F7 monoclonal | sc-59737 | Unconjugated | Mouse | Santa Cruz | 1:200 |
| Rabbit | N/A | A-11008 | Alexa Fluor 488 | Goat | Life Technologies | 1:800 |
| Rabbit | N/A | A-11036 | Alexa Fluor 568 | Goat | Life Technologies | 1:800 |
| Goat | N/A | A-11055 | Alexa Fluor 488 | Donkey | Life Technologies | 1:800 |

| Antigen | Clone | Catalog # | Conjugate | Host | Supplier | Dilution |
|---|---|---|---|---|---|---|
| Goat | N/A | A-11058 | Alexa Fluor 594 | Donkey | Life Technologies | 1:800 |
| Mouse | N/A | A-11031 | Alexa Fluor 568 | Goat | Life Technologies | 1:800 |

**Supplementary Table 2.** The primers used for qPCR.

| Gene | Forward primer | Reverse primer | Length [bp] |
|---|---|---|---|
| ***Actb*** | GGCTGTATTCCCCTCCATCG | CCAGTTGGTAACAATGCCATGT | 154 |
| ***Aqp5*** | AGAAGGAGGTGTGTTCAGTTGC | GCCAGAGTAATGGCCGGAT | 220 |
| ***Axin2*** | TGACTCTCCTTCCAGATCCCA | TGCCCACACTAGGCTGACA | 105 |
| ***Calca*** | GAGGGCTCTAGCTTGGACAG | AAGGTGTGAAACTTGTTGAGGT | 101 |
| ***Cftr*** | CCCTTCGGCGATGCTTTTTC | AAGCCTATGCCAAGGTAAATGG | 166 |
| ***Cyp2f2*** | GGACCCAAACCTCTCCCAATC | CCGTGAACACCGACCCATAC | 106 |
| ***Eef1g*** | TTCCTGCCGGCAAGGTTCCA | TGCCGCCTCTGGCGTACTTC | 119 |
| ***Epcam*** | GCGGCTCAGAGAGACTGTG | CCAAGCATTTAGACGCCAGTTT | 139 |
| ***Foxi1*** | CCTCTCCACCATGACAGCAT | TCCCATGGCTACTGAGGTTG | 155 |
| ***Foxj1*** | CCCTGACGACGTGGACTATG | GCCGACAGAGTGATCTTGGT | 114 |
| ***Krt5*** | CTCTGTCGTTACAAACAGTG | CTTAGCCCGCTACCCAAACC | 159 |
| ***Krt8*** | CAAGGTGGAACTAGAGTCCCG | CTCGTACTGGGCACGAACTTC | 187 |
| ***Muc5ac*** | GTGGTTTGACACTGACTTCCC | CTCCTCTCGGTGACAGAGTCT | 103 |
| ***Muc5b*** | TCCCTAGCATGAGCGCCTTA | CCACGACGCAGTTGGATGTT | 178 |
| ***Pdpn*** | ACCGTGCCAGTGTTGTTCTG | AGCACCTGTGGTTGTTATTTTGT | 159 |
| ***Scgb1a1*** | ATGAAGATCGCCATCACAATCAC | GGATGCCACATAACCAGACTCT | 135 |
| ***Sftpa1*** | GAGGAGCTTCAGACTGCACTC | AGACTTTATCCCCCACTGACAG | 103 |
| ***Sftpb*** | CTGCTTCCTACCCTCTGCTG | CTTGGCACAGGTCATTAGCTC | 175 |
| ***Sftpc*** | ATGGACATGAGTAGCAAAGAGGT | CACGATGAGAAGGCGTTTGAG | 117 |
